# Changes in spinal cord hemodynamic reflect modulation of spinal network with different parameters of epidural stimulation

**DOI:** 10.1101/833202

**Authors:** Shanshan Tang, Carlos A. Cuellar, Pengfei Song, Riazul Islam, Chengwu Huang, Hai Wen, Bruce E. Knudsen, Ping Gong, U-Wai Lok, Shigao Chen, Igor A. Lavrov

## Abstract

In this study functional ultrasound (fUS) imaging has been implemented to explore the local hemodynamic response induced by electrical epidural stimulation and to study real-time *in vivo* functional changes of the spinal cord, taking advantage of the superior spatiotemporal resolution provided by fUS. By quantifying the hemodynamic and electromyographic response features, we tested the hypothesis that the transient hemodynamic response of the spinal cord to electrical epidural stimulation could reflect modulation of the spinal circuitry and accordingly respond to the changes in parameters of electrical stimulation. The results of this study for the first time demonstrate that the hemodynamic response to electrical stimulation could reflect functional organization of the spinal cord. Response in the dorsal areas to epidural stimulation was significantly higher and faster compared to the response in ventral spinal cord. Positive relation between the hemodynamic and the EMG responses was observed at the lower frequencies of epidural stimulation (20 and 40 Hz), which according to our previous findings can facilitate spinal circuitry after spinal cord injury, compared to higher frequencies (200 and 500 Hz). These findings suggest that different mechanisms could be involved in spinal cord hemodynamic changes during different parameters of electrical stimulation and for the first time provide the evidence that functional organization of the spinal cord circuitry could be related to specific organization of spinal cord vasculature and hemodynamic.

**Significance Statement:** Electrical epidural stimulation (EES) has been successfully applied to control chronic refractory pain and was evolved to alleviate motor impairment after spinal cord injury, in Parkinson’s disease, and other neurological conditions. The mechanisms underlying the EES remain unclear, and current methods for monitoring EES are limited in sensitivity and spatiotemporal resolutions to evaluate functional changes in response to EES. We tested the hypothesis that the transient hemodynamic response of the spinal cord to EES could reflect modulation of the spinal cord circuitry and accordingly respond to the changes in parameters of EES. The proposed methodology opens a new direction for quantitative evaluation of the spinal cord hemodynamic in understanding the mechanisms of spinal cord functional organization and effect of neuromodulation.

## Introduction

Electrical epidural stimulation (EES) of the spinal cord has been implemented earlier to alleviate chronic pain and in the last decade, clinical trials have shown that EES can restore volitional, motor function in patients suffering spinal cord injury to some extent (references). Interestingly, participants on those trials have also experienced improvements in autonomic function, namely cardiovascular, sexual, and voiding, among others. Additionally, reports on other neurological conditions have shown benefits from EES [1–10]. Regardless the wide application of this technique, there is still limited understanding of the mechanisms of EES effect on spinal circuitry.

Computational models [1, 11, 12] and available *in vivo* and *in vitro* electrophysiological assessments [13–17] have shed some light on the potential components of neural circuits activated by EES, although, this information is limited and cannot provide deep insight into neurophysiologic mechanisms of EES effect. This is largely due to the absence of sensitive and reliable tools that could provide critical information about spatial-temporal activation of spinal circuits. For instance, current approaches aim to target the spinal circuitry by using neuromodulation with electrical, magnetic, or pharmacological influence, while using electrophysiological assessment for output measuring i.e. electromyography (EMG). This wide used recording technique provides one dimensional signal reflecting the electrical activity produced by skeletal muscles as a reflection of activation level of neural circuit and although useful, is susceptible to the noise generated by movements during breathing or during muscle contraction caused by electrical stimulation.

Functional neuroimaging techniques, such as fMRI, PET, and MEG are successful in mapping functional organization of the brain [20–22]; however, these methods have not yet been very effective in mapping organization of spinal cord due to challenging location and anatomy of the spinal cord. Recently, functional ultrasound imaging (fUS), a sensitive imaging modality with the use of plane-wave ultrasound illuminations at high frame rate, has achieved valuable spatiotemporal resolution to capture the transient hemodynamic response [23–26]. As an emerging novel technique, fUS has been successfully employed to investigate the transient changes in blood volume in the whole brain on animal models with better spatiotemporal resolution compared to the aforementioned functional brain imaging modalities [23, 24].

Previously we reported hemodynamic changes in the spinal cord during EES in two animal models as monitored by fUS. We also described that fUS offers a better sensitivity in mapping and quantifying hemodynamics in the spinal cord and quantifying the spinal cord activity during subthreshold-to-threshold motor responses produced by EES compared to electrophysiological assessment [25]. These results suggests that fUS can provide assessment of spinal circuitry at subthreshold level of stimulation, when only low threshold afferents are active, which significantly improves capacity to evaluate the spinal circuitry organization *in vivo*.

In this study we extend our previous observations and hypothesized that the spinal cord hemodynamic response to EES reflects a transient modulation of spinal cord circuitry and, accordingly, responds to the changes in parameters of electrical stimulation. In order to test this hypothesis, we evaluated the spinal cord hemodynamic response to the different frequencies, electrodes configurations, as well as different voltage intensities; and quantified the features of hemodynamic response, such as peak increase in blood volume change, the response rate, and the total amount of blood volume change over the time. With this study, we for the first time provide evidence that functional organization of the spinal cord circuitry is related to organization of spinal cord hemodynamic as revealed by implementation of a novel fUS approach.

## Materials and Methods

### Animal preparation procedure

Experiment procedures were approved by the Mayo Clinic Institutional Animal Care and Use Committee. The National Institutes of Health Guidelines for Animal Research (Guide for the Care and Use of Laboratory Animals) were observed rigorously. Animals were kept in controlled environment (21°C, 45% humidity) on a 12-h light/dark cycle with ad libitum access to water and food.

Eight male Sprague-Dawley rats (325-350 gr) were anesthetized with 1.5%-3% isoflurane. The spinal cord was exposed by removing lamina T13-L2 (corresponding approximately to L2 and S1 segments of the spinal cord). Teflon coated stainless steel wires were placed at T13 and L2 and then sutured on dura. A 0.5mm notch facing the spinal cord was made on the wires, serving as the stimulating electrode. The spine was fixed to minimize the breathing motion with a custom-made frame composed of a clamp holding the Th12 spinous process and two pieces retracting back muscles on both sides. In addition, the pelvic girdle was held with two rods secured over the coxal bones. Dorsal skin flaps were attached around the frame to form a pool facilitating transducer positioning. Warm saline solution (1.5 ml) was administered S.C. every 2 h. At the end of experiment, animals were euthanized using pentobarbital (150 mg/kg I.P.).

### Experiment Setup

A Verasonics Vantage ultrasound system (Verasonics Inc., Kirkland, WA) and a Verasonic high frequency linear array transducer L22-14v (Verasonics Inc., Kirkland, WA) were used in this study to provide transmission waveforms with 15.625 MHz center frequency and 67% bandwidth. The transducer was placed between the T13 and L2 vertebrae with an ultrasound imaging field of view (FOV) align with the spinal cord and intersect with the central canal, as shown in Fig. 1(a) and 1(b). The transducer was fixed thoroughly throughout the experiment. Mineral oil was used as acoustic coupling between the spinal cord and ultrasound transducer.

**Fig. 1.**
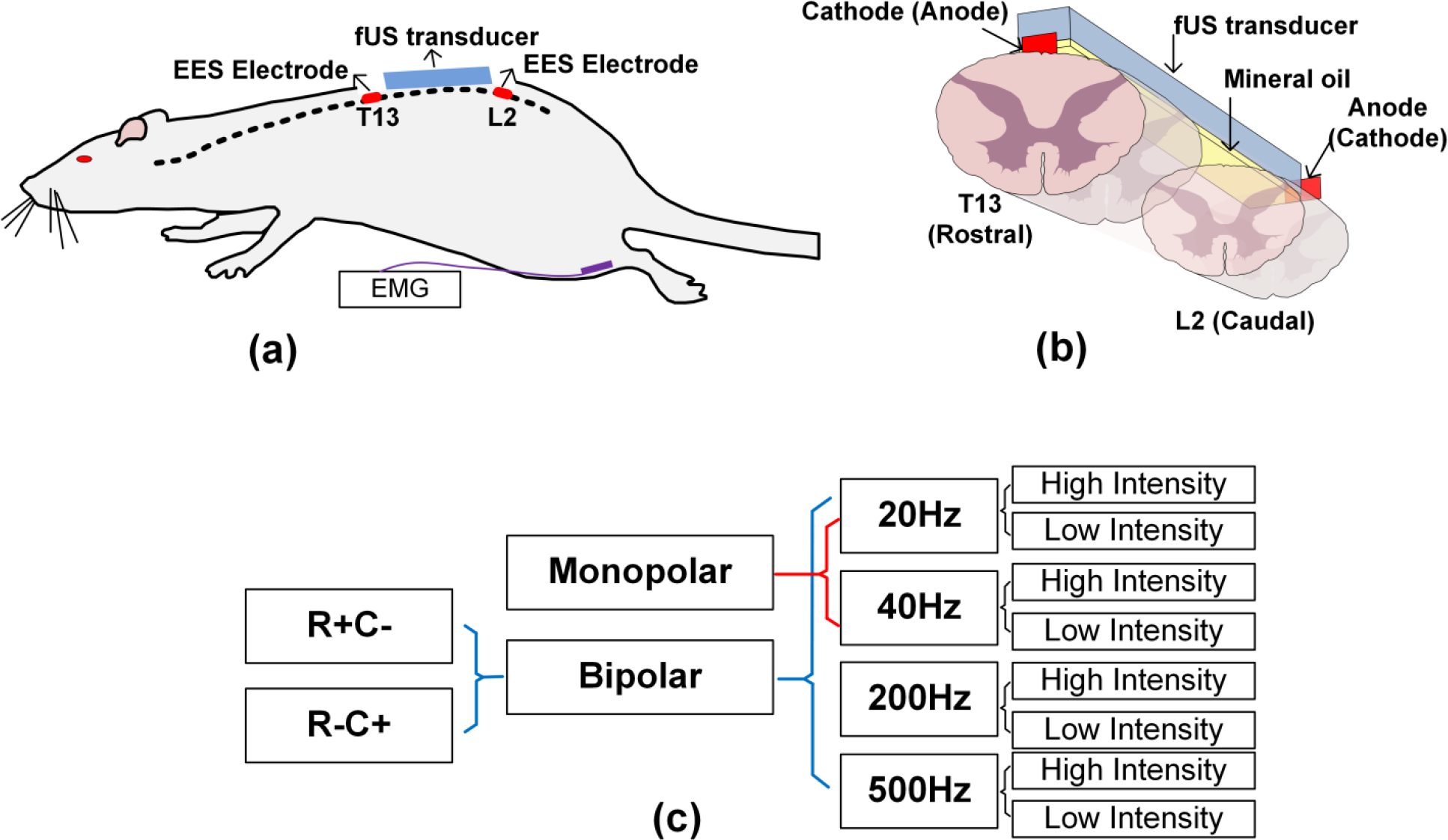
Experiment setup and EES parameters and electrode configurations. (a) and (b) illustrates the positioning of ultrasound transducer, spinal cord electrodes, and the EMG electrodes. (c) Detailed EES configurations. R+C− represents the anode is placed at spinal cord T13 while the cathode is placed at spinal cord L2.R−C+ represents the opposite placement (R for rostral and C for caudal accordingly).

To increase the imaging signal-to-noise-ratio (SNR), ultrafast compounding plane wave imaging sequence was used with five steered plane waves transmitted with each steering angle (−4°, −2°, 0°, 2°, 4°) for three repetitions [27]. With a pulse repetition frequency (PRF) of 28.6 kHz, it took 525 μs to complete the 15 transmissions. A no operation period of 1475 μs was added to the end of the 15 transmissions, to achieve a post-compounding PRF of 500 Hz. The total data recording time was 120 s with 200 ensembles in each second.

In the rat experiment, the stimulation protocol was set as 30 s baseline measurements, 20 s EES measurements, and 70 s recovery measurements. Five trials were repeated with the exactly same EES configurations.

### ESS parameters and electrodes configurations

EES was delivered on the spinal cord of a rat at vertebrae levels T13 and L2. Different stimulation polarities, including monopolar and bipolar stimulations, were used. Monopolar stimulation was applied with the cathode (−) placed on the T13 spinal cord and anode (+) placed in rat’s back muscle. In bipolar stimulation, the anode and cathode electrodes were either placed on T13 and caudal L2 spinal cord segments (R+C−, “R” for rostral and “C” for caudal accordingly), or in L2 and T13 spinal cord segments (R-C+), respectively. Both low frequency ranges of EES (20 Hz and 40 Hz) and high frequency EES (200 Hz and 500 Hz) were used in this study and were chosen based on our previous findings with significantly variations of stepping performance with EES applied with these two frequencies in spinal rats [28–33]. Both sub-threshold intensity and supra-threshold intensity electrical stimulation were tested. Due to the difference in motor threshold across the animals, stimulation voltages were carefully selected based on real-time evaluation of EMG signal and gradual increase and decrease of intensity of EES. The supra-threshold intensity was determined by increasing the voltage to a maximum level at which stimulation produced a strong contraction, without tremble affecting the fUS imaging. The sub-threshold intensity was determined by decreasing the voltage to a subthreshold level where EMG signal was undistinguished from noise. A summary for the detailed EES parameters and electrode configurations can be found in Fig. 1(c).

### Data Processing

After coherent compounding, B-mode ultrasound images were processed with phase correlation based sub-pixel motion registration [34] and singular-value-decomposition (SVD) based cluttering filtering [35, 36] to accumulate one frame of Power Doppler (PD) image in each second.

Ultrasound PD signal was measured at each imaging pixel to reflect the local blood volume. The spinal cord hemodynamic response to EES was evaluated by the spinal cord blood volume change (ΔSCBV), which defines the PD signal intensity variation along with the EES process in the unit of percentage,

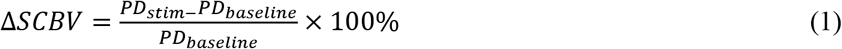

A ΔSCBV response curve under an EES configuration was obtained by averaging the ΔSCBV response curve of each pixel inside a selected region of interest (ROI) and averaging all 5 repeated trials under this EES configuration. The ROIs used in this study include the whole ultrasound FOV, dorsal ROI, ventral ROI, rostral ROI, and caudal ROI. From a ΔSCBV curve, quantitative parameters, including the peak response, ascending slope of the response curve (response rate), and area under the curve (AUC) were measured, as illustrated in Fig. 2(a). Detailed data processing approach could be found in our previous study [25].

**Fig. 2.**
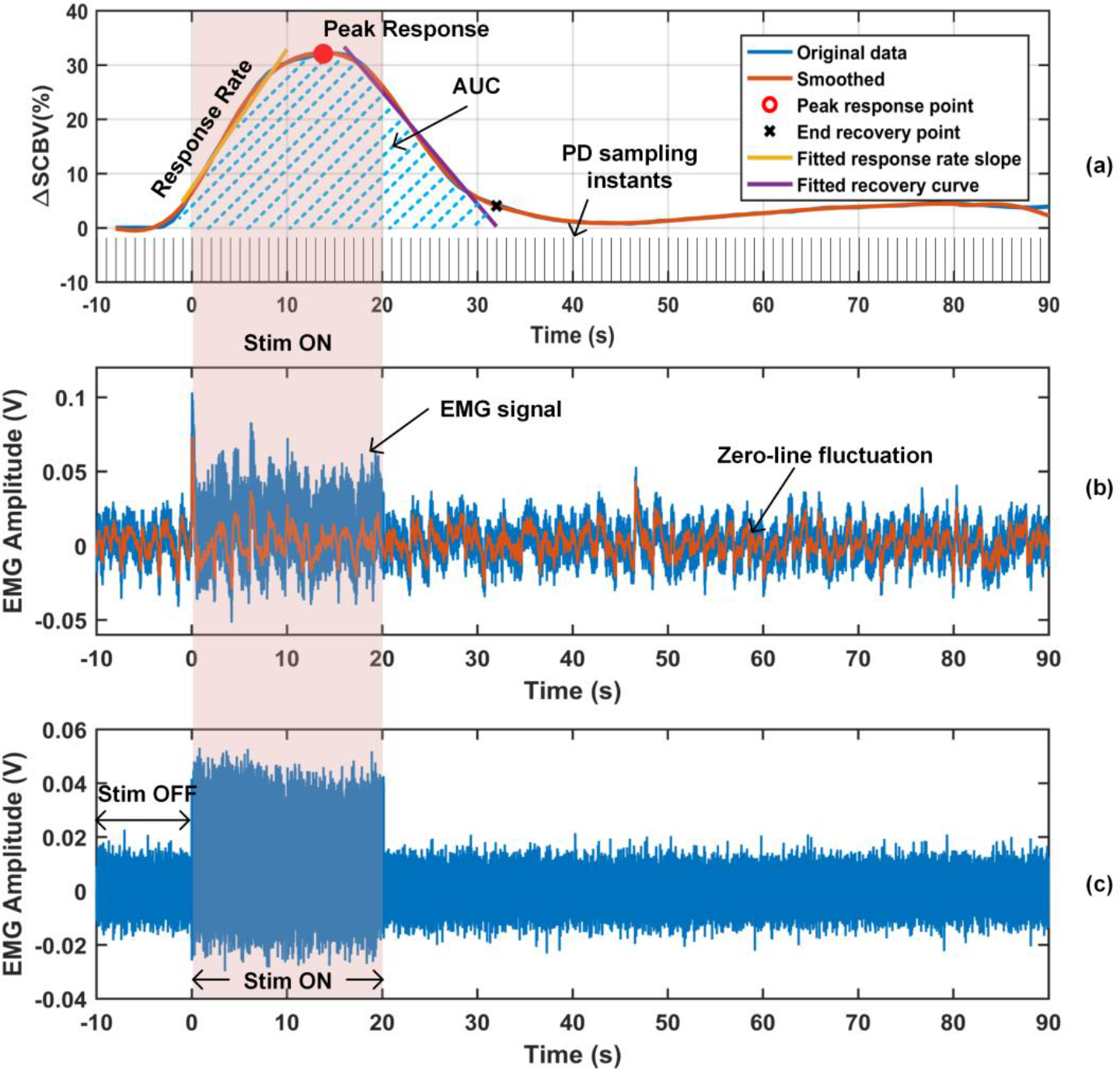
ΔSCBV response curve and EMG signal. (a) Parameters including peak response, response rate, and area under the curve (AUC), were measured from a ΔSCBV curve. (b) EMG signal with zero-line fluctuation. (c) EMG signal with the zero-line fluctuation removed.

EMG signals from the tibialis anterior and gastrocnemius hind limb muscles were recorded simultaneously in experiments. The fluctuated zero-line of each EMG signal (Fig. 2(b)) was first extracted and then subtracted from the EMG signal (Fig. 2(c)). Then, the root mean square (RMS) of the EMG signal during EES and the RMS of background activity before EES were measured and ΔEMG was calculated in the unit of percentage,

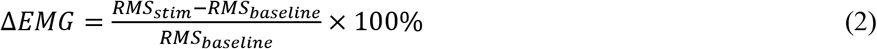

where *RMS*_*baseline*_ and *RMS*_*stim*_ indicates the RMS of the signal during “Stim OFF” and “Stim ON” in Fig. 5(c). ΔEMG was averaged for the 5 repeated trials for each of the four hind limb muscles where EMG signals were recorded from. The largest ΔEMG among the four was selected to represent the neuromuscular electrophysiological change under the given EES.

## Results

### A. fUS is a sensitive tool in quantifying the subthreshold-to-threshold motor activation response to EES

During initial testing, we found that fUS offers valuable sensitivity in quantifying the hemodynamic response in the spinal cord during subthreshold EES, this is even when the amplitude of motor responses are not distinguishable from electrical noise in the EMG recording. A typical hemodynamic response to the sub-threshold EES intensity is shown in Fig. 3(a) and Supplemental video 1, where the ultrasound power Doppler (PD) images were color coded with the measured blood volume change, ΔSCBV. The corresponding ΔSCBV curve is shown in Fig. 3(b), from which an obvious hemodynamic response to the EES can be observed primary in the dorsal part of the spinal cord. At the same time, no statistically significant changes in the EMG signal were observed due to the sub-threshold intensity of EES (Fig. 3(c)).

**Fig. 3.**
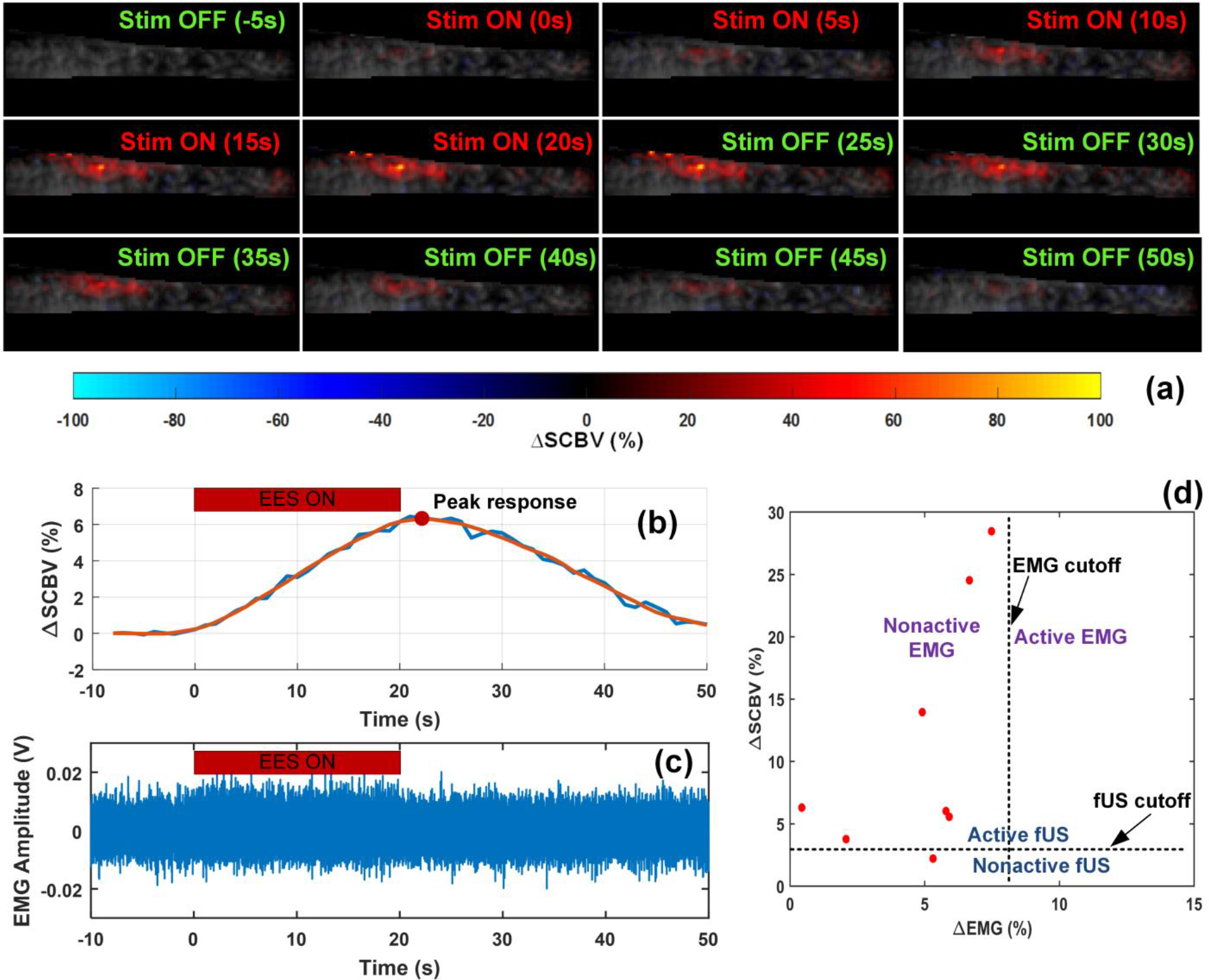
fUS is a sensitive technique in monitoring the hemodynamic response of the spinal cord even during subthreshold EES intensities (i.e. silent EMG). (a) Spinal cord blood volume change (ΔSCBV) color coded on the Power Doppler (PD) images at several point times. A movie is available in the Supplemental video 1. (b) and (c) are ΔSCBV curve and ΔEMG signal for a case with sub-threshold EES and silent EMG. (d) ΔSCBV peak response vs ΔEMG for the 8 cases with silent EMG signals.

EES voltage intensity inducing subthreshold level of motor responses varied across animals and for sub-threshold motor potentials evoked by EES, the intensity was determined by gradually decreasing the voltage from a readily observable EMG response to a subthreshold level with no obvious EMG signal. Among EMG recordings with sub-threshold EES, we applied an offline processing, where an EMG signal was considered as “silent EMG” only when its root mean square (RMS) increase of EMG signal to the baseline, ΔEMG, is lower than 8% (see Supplemental Fig. S1(a)). During experiments we collected 25 cases (a case means data collected from a given animal with a given EES parameter and electrode configuration) with sub-threshold EES, among which 8 cases met the criteria and considered as silent EMG. In these 8 cases EES was applied at sub-threshold level with frequencies of 20 Hz, 40 Hz, and 200 Hz, and also with monopolar electrode configurations, with bipolar R−C+ configuration, and with bipolar R+C− configuration. Similarly, a ΔSCBV with a peak response increase greater than 3% was considered reliable in hemodynamic response parameter measurements (see Supplemental Fig. S1(b)-1(c) and Supplemental video 2 and video 3). The peak response of the ΔSCBV curve was measured for the aforementioned 8 cases, where silent EMG signals were observed. Seven cases of the 8 exhibited obvious responses in fUS imaging with a peak response above the preset cutoff, as shown in Fig. 3(d), suggesting that evaluation of spinal cord hemodynamic with fUS is more sensitive compared to EMG monitoring during EES.

### B. Spinal cord hemodynamic response to EES is spatially dependent

To evaluate the effect of EES-induced changes in blood volume in different regions of the spinal cord, ΔSCBV response curves and their response parameters were measured in selected spinal cord regions under various EES configurations. Figure 4(a) depicts typical color-coded hemodynamic response images at their peak blood volume increasing induced by EES with distinct parameters and electrode configurations (case number, n=4). The ultrasound FOV is divided in dorsal and ventral regions by the white dashed lines and to rostral and caudal regions by the green dashed lines.

**Fig. 4.**
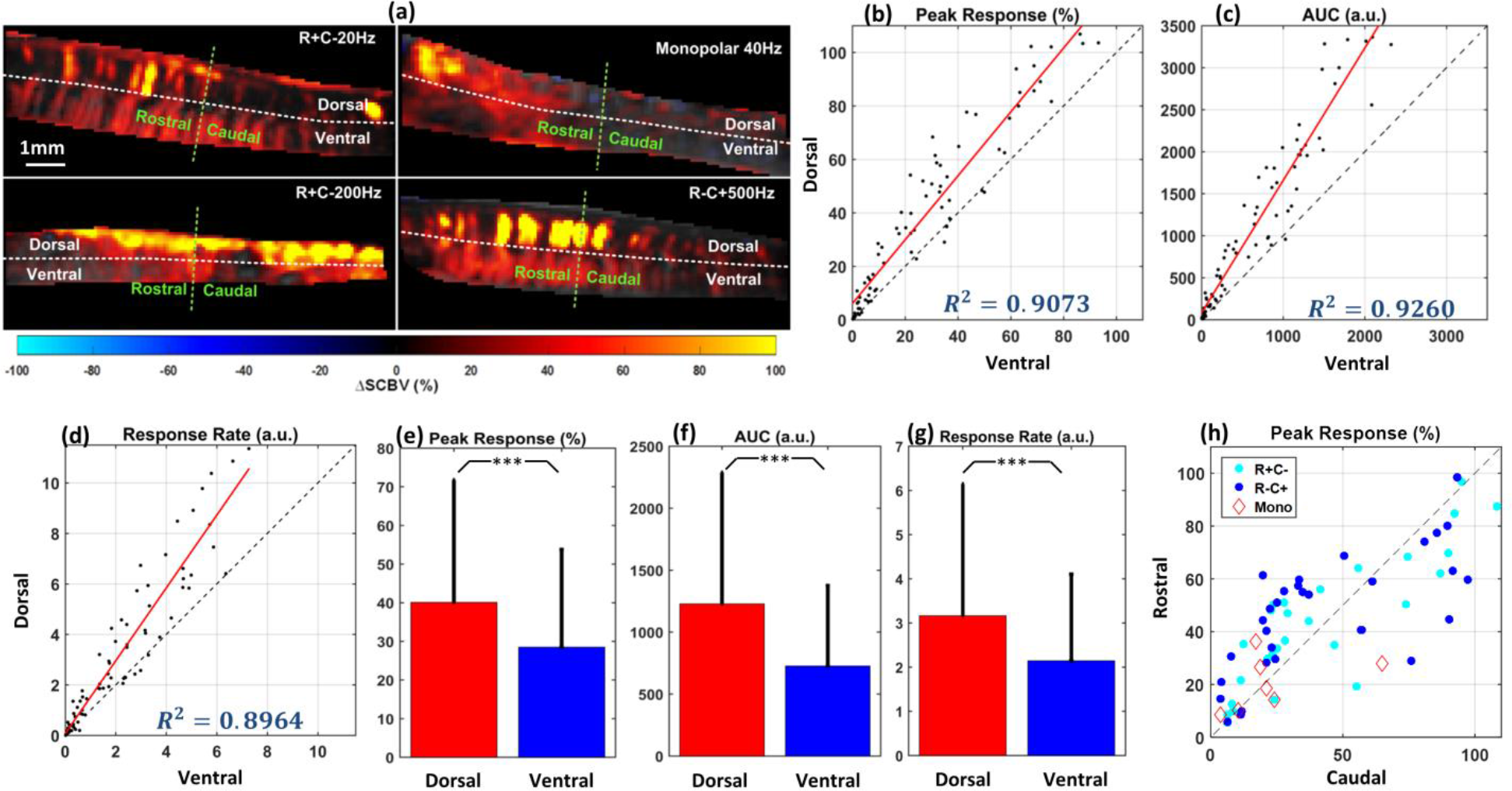
Spinal cord hemodynamic response due to EES is spatially dependent. (a) Representative examples of hemodynamic response images with different EES parameters and electrode configurations. The four fUS images were collected from supra-threshold intensities, different stimulation frequencies (20 Hz, 40 Hz, 200 Hz and 500 Hz) and electrode configurations (monopolar, bipolar R+C−, and bipolar R−C+). Ultrasound FOV is divided in dorsal and ventral regions by the white dashed line, and to the rostral and caudal regions with the green dashed line. (b) Peak response of blood volume change in dorsal region vs ventral region (case number, n=78). (c) Total blood volume change over the time (Area under the curve, AUC) for dorsal and ventral regions (case number, n=78). (d) Response rate for dorsal and ventral regions (case number, n=78). (e)-(g) Bar plots of the peak response, AUC and response rate for dorsal and ventral regions. Hemodynamics of the dorsal region of the spinal cord responded significantly higher and quicker to EES compared to ventral regions, regardless of the stimulation parameter and configuration. (h) Peak response for rostral and caudal regions (case number, n=78). No significant difference between rostral and caudal region was found for either configuration of electrodes. Each data point in (b)-(d) and (h) represents the peak blood volume change in dorsal/ventral or rostral/caudal regions from the same rat with the same stimulation parameter and configuration. Results in (b)-(h) includes all tested EES parameters and electrode configurations (EES intensity includes sub-threshold and supra-threshold; EES frequency includes 20 Hz, 40 Hz, 200 Hz and 500 Hz; EES electrode configuration includes monopolar, bipolar R+C−, and bipolar R−C+). *** p<0.001.

#### B1. Dorsal vs. Ventral

Figure 4(b) and 4(c) show the peak response and AUC in the dorsal and ventral regions of the spinal cord, while Fig. 4(e) and 4(f) are the corresponding bar plots. Data shown in Fig. 4(b) and 4(c), also in Fig. 4(e) and 4(f), includes 8 rats’ responses induced by EES with sub-threshold and supra-threshold intensities, 20 Hz, 40 Hz, 200 Hz, and 500 Hz stimulation frequencies, and monopolar, bipolar R+C−, and bipolar R−C+ electrode configurations (case number, n=78). Each dot on Fig. 4(b) and 4(c) represents the dorsal vs ventral hemodynamic responses from the same rat with the same EES parameter and electrode configuration. Significant differences (paired t-test, *p<0.001*) were observed in both peak response and AUC between dorsal and ventral spinal cord regions, as illustrated in Fig. 4(e) and 4(f). A significant difference in response rate was also observed between dorsal and ventral regions (paired t-test, *p<0.001*) at almost all tested EES configurations, as shown in Fig. 4(d) and 4(g) (case number, n=78). Overall, the dorsal region of the spinal cord responded with a higher magnitude and a quicker response rate to the EES compared to the ventral region, independently to the tested stimulation configurations.

#### B2. Rostral vs. Caudal

To evaluate the role of electrode configurations in spinal cord hemodynamic response, rostral vs. caudal peak responses of ΔSCBV recorded with monopolar stimulation, bipolar R+C−, and bipolar R−C+ stimulation were compared (Fig. 4(h)). Rostral and caudal regions were defined with the middle line perpendicular to the spinal cord FOV, which was between the T13 and L2 vertebrae. Each dot in Fig. 4(h) represents the rostral versus caudal hemodynamic responses from the same rat with the same EES parameter and electrode configuration. Different colors represent cases with different electrode configurations. Results of Fig. 4(h) were recorded from 8 rats with sub-threshold and supra-threshold intensities, 20 Hz, 40 Hz, 200 Hz, and 500 Hz stimulation frequencies, and monopolar and bipolar electrode configurations (case number, n=78). No significant difference between rostral and caudal region was found for either configuration, indicating that the basic configurations of epidural electrodes have no significant influence on the spatial distribution of spinal cord hemodynamic response. At the same time, we observed that the rostral-caudal response pattern varied from animal to animal. Among the 8 tested rats, 5 showed significantly higher ΔSCBV response in the rostral region compared to the caudal region (paired t-test, *p<0.001*), while the rest of three rats showed significantly higher ΔSCBV response in caudal region compare to the rostral region (*p<0.001*), regardless of the electrode configuration (results not shown).

### C. Spinal cord hemodynamic response to EES is frequency dependent

ΔSCBV peak responses with stimulation frequencies of 20 Hz and 40 Hz compared to stimulation frequencies of 200 Hz and 500 Hz were plotted as a function of ΔEMG, as shown in Fig. 5(a)-(b). Data presented on Fig. 5 were collected from 5 rats with bipolar EES configuration (R+C− and R−C+) and supra-threshold intensity (case number, n=14 and n=13 for Fig. 5(a) and 5(b) respectively). At low stimulation frequencies (20 and 40 Hz), ΔSCBV was positively correlated to ΔEMG (Fig. 5(a)), suggesting that at those frequencies, the spinal cord neuronal activity is related to the hemodynamic response. However, no positive relationship was found in either 200 Hz or 500 Hz (Fig. 5(b)). Figure 5(c) shows the response rate between 40 Hz and 200 Hz EES, chosen based on our previous findings [28–33]. Each data point in Fig. 5(c) represents the response rates measured from the two frequencies, but the same rat and the same electrode configuration (case number, n=13). Comparison of spinal cord hemodynamic response during 40 Hz and 200 Hz indicates that the spinal cord blood flow increases at 40 Hz stimulation significantly faster than at 200 Hz (paired t-test, *p<0.001*) (Fig. 3(c)).

**Fig. 5.**
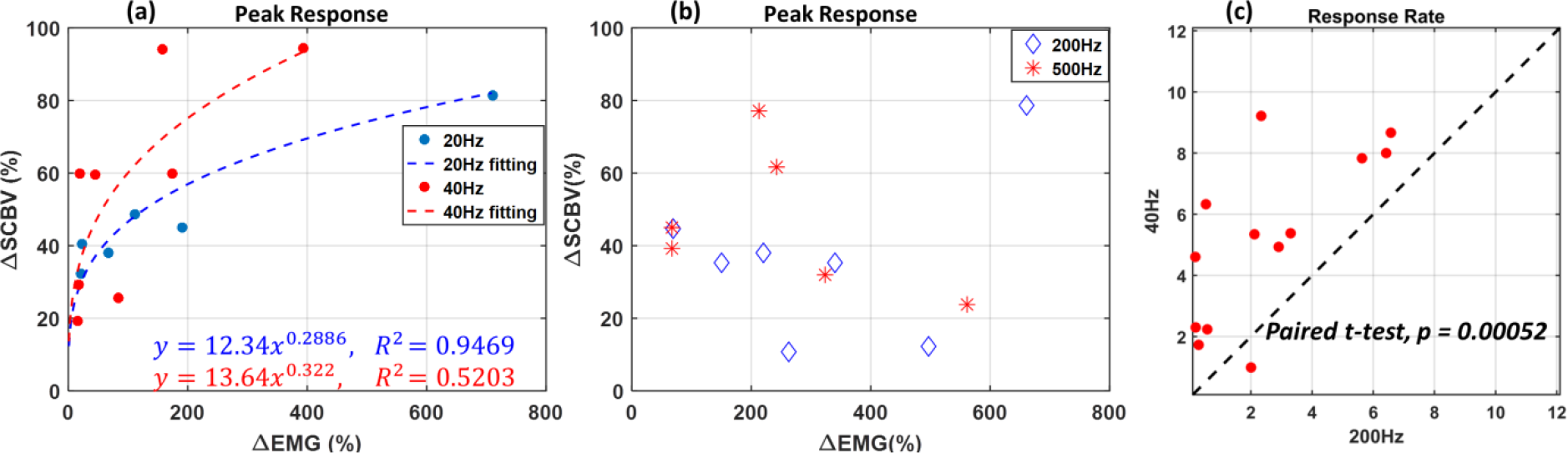
EES produce frequency dependent hemodynamic response within spinal cord. (a) Increase in the hemodynamic response was observed with stimulation frequencies of 20 and 40 Hz (case number, n=14), (b) but not with higher stimulation frequencies of 200 and 500 Hz (case number, n=13). Data presented in (a) and (b) are with electrode configurations including bipolar R+C− and bipolar R−C+, and supra-threshold intensity. (c) Response rate of the spinal cord to low (40 Hz) and high (200 Hz) stimulation frequencies chosen based on our previous findings [28–33] (case number, n=13). Each dot represents the response rate measured from the two frequencies but the same rat and the same electrode configuration. Data presented in (c) are with supra-threshold intensity and electrode configuration including bipolar R+C− and bipolar R−C+.

## Discussion

The results of this study for the first time show differential hemodynamic responses in dorsal vs. ventral spinal cord during EES evidenced by fUS. We found specific features related to parameters of stimulation: (1) even at low stimulation intensities (subthreshold to EMG response), EES induced clear hemodynamic changes in the dorsal, but not ventral spinal cord, even though the EMG output was silent, which may indicate on critical role of the dorsal horn neurons cumulatively processing a greater amount of the sensory information from dorsal roots afferents compared to low threshold muscle afferent activating the motoneurons and premotor interneurons; (2) significantly higher and faster hemodynamic responses in dorsal regions compared to the ventral spinal cord, and (3) significant differences in spinal cord hemodynamic responses at low and high stimulation frequencies, with a positive relation between the hemodynamic and the EMG response observed at lower frequencies (20 Hz and 40 Hz), but not at higher frequencies (200 Hz and 500 Hz). These results support our initial observation that evaluation of spinal cord hemodynamic provides significantly higher sensitivity compared to *in vivo* electrophysiological assessment, such as EMG monitoring.

Motor threshold responses during EES were determined based on EMG amplitude, which was higher than background level of activity, so results with ΔEMG lower than a given cutoff were considered as silent EMG data (Supplemental Fig. S1(a)). As we demonstrate in this study, EES facilitated hemodynamic response primary in dorsal regions of spinal cord, regardless of parameters and electrodes configurations, which further support high sensitivity of fUS in evaluation of local changes in the spinal cord during EES. This further suggests that evaluation of spinal hemodynamic changes may help in optimizing electrodes location and expanding our understanding of the mechanisms of spinal cord neuromodulation and spinal cord functional organization.

The observed hemodynamics changes are likely correlated with spinal cord functional neuroanatomy and particularly with specific organization of vasculature for the dorsal and ventral spinal cord. The ventral part of the spinal cord has predominantly segmental organization, arranged as a string of beads, while the dorsal regions are usually arranged as one solid column, although varying in thickness at different levels [37–39]. This specific anatomical organization of the spinal cord vasculature and unique distribution of dorsal and ventral spinal arteries, to some extent divides the spinal cord into two functionally different areas of hemodynamics [40, 41], which could to some extent explain different response of the dorsal vs. ventral parts to EES. Although, this unique anatomical difference in organization of the spinal cord vasculature has been known for years, the results of this study for the first time show that similar difference exists in functional organization of the dorsal vs. ventral parts of the spinal cord.

Interestingly, in this study the orientation of electric field adjusted by variations in polarity of stimulating electrodes did not induce significant changes in spinal cord hemodynamics, as shown in Fig. 2(h), although the rostral-caudal distribution of hemodynamic response was variable between tested animals, which could be due to the two-dimension-sectional nature of fUS FOV, which images one longitudinal cross-section of the spinal cord, considering the 3mm spinal cord diameter is thicker than the elevational resolution of the high-frequency transducer used in the study.

The results of this study suggest that fUS has relatively high spatial and temporal resolution in monitoring the localized hemodynamic response at different spinal cord segments [25]. Although other functional imaging modalities, such as PET and fMRI, have their distinct advantages, such as high sensitivity and excellent imaging contrast and imaging depth, their spatial and/or temporal resolution are far below what is required for evaluation of the spinal cord functional changes, particularly during neuromodulation with EES. For instance, the imaging FOV of the presented fUS images was 9.9 mm ×12.8 mm with a spatial resolution of 98.56 μm (the wavelength of ultrasound pulses, as well as the pixel size of fUS image) and a temporal resolution of 500 Hz. In contrast, typical spatial resolution of fMRI is 1.5 mm −3 mm. fMRI with a 7 Tesla machine reaches a spatial resolution of 750 μm by sacrificing the tSNR and detecting sensitivity [42, 43]. In addition, the size of MR or PET machine can be prohibitive for an intraoperative monitoring. Although functional neuroimaging techniques including EEG and MEG have very high temporal resolution, they could not be applied to study the stimulation related hemodynamic response, where fUS is able to complement.

Factors such as temperature change, autonomic regulation, strenuous activities, etc., may also induce changes in spinal cord blood flow, which can be detected by fUS. Our previous study [25] demonstrated that during background fUS recording of the spinal cord without electrical stimulation no increase of blood flow signal was observed across multiple trials.

The results of this study for the first time emphasize a coupling between the spinal cord hemodynamics and spinal circuits activated by EES at low frequencies (20 Hz – 40 Hz) compared to high frequencies (200 Hz – 500 Hz) and support the hypothesis that the modulation in parameters of EES may activate different mechanisms and/or different components of the spinal circuitry. Similar results were presented earlier [44] where the cerebral blood flow in rat somatosensory cortex was found to be highly correlated to the field potentials of electrical stimulation on the infraorbital nerve at low frequencies (<2 Hz) and uncoupling at stimulation frequencies of 2– 5 Hz. Although the frequency range in both studies is different that could be due to the different parts of the nervous system and different neural organization. A coupling between the spinal cord hemodynamics and the neuronal activity facilitated at frequency range 20 Hz – 40 Hz is supported by our early findings where facilitation of the spinal circuitry to maintain optimal stepping pattern was successfully achieved with epidural stimulation at the similar frequency range [28–33, 45]. Although current results may not be sufficient to draw a certain conclusion regarding how particular frequency modulates the spinal cord circuitry, the mechanisms responsible for this effect can be evaluated in future studies by selective stimulation of spinal structures and peripheral nerves or by comparing the effect of EES with pharmacological activation or inhibition of the different components of spinal network.

The results of this study also support a great potential of fUS as a research tools to study spinal cord and as a potential diagnostic tool with several critical clinical applications. Currently, there is no available technique that could evaluate functional changes in spinal cord in real time and fUS technique with miniaturized transducer size can be implemented for real-time and long-term monitoring. Spinal cord fUS could also help in evaluation of hemodynamic response during electrode implantation to optimize leads location for neuromodulation therapies and for intraoperative monitoring, which cannot be achieved by other functional imaging modality, such as fMRI or PET. fUS may also help to reveal important information about spinal cord circuitry activated during pharmacological interventions and different modalities of neuromodulation to tune dosage or stimulation parameters for more specific delivery of these therapies. The findings that subthreshold epidural stimulation can selectively increase hemodynamic in the dorsal areas of the spinal cord could also lead to development of novel combined pharmacological and neuromodulation treatments, where local pharmacological effect could be further enhanced by activation of local hemodynamic in target area. Spinal cord fUS can be also utilized to investigate the mechanism of pain development and effect of spinal cord neuromodulation, by characterizing associated neurovascular coupling [46–48]. Other potential applications could include evaluation of spinal cord ischemia and revascularization following SCI, local tumor growth, and others [49–51].

## Conclusions

In summary, results of this study indicate that (1) the dorsal regions of the spinal cord have significantly higher and faster blood volume changes compared to ventral spinal cord region during EES, regardless of parameters and electrodes configurations; (2) spinal cord hemodynamic changes are more responsive to the low frequencies (20-40 Hz) of stimulation, at which ΔEMG and ΔSCBV were positively related, that was not observed at high EES frequencies (200-500 Hz), suggesting the frequency-dependent mechanism of spinal cord hemodynamic response; (3) hemodynamic changes recorded with fUS provide more sensitive approach for evaluation of the spinal cord response to neuromodulation compare to currently available *in vivo* electrophysiological techniques. This study for the first time explores the role of spinal cord hemodynamic in effect of EES and suggests that functional organization of spinal hemodynamics could be coupled with spinal neural circuitry. These findings support future development of neuromodulation techniques where the stimulation parameters, electrodes location, and the orientation of the electric field could be precisely tuned based on real-time evaluation of the local changes in spinal cord hemodynamic. Although the mechanisms of presented here findings need further investigations, spinal cord fUS opens a new direction for quantitative evaluation of spinal cord hemodynamic and understanding the spinal cord functional organization and the mechanisms of spinal cord neuromodulation.

## Acknowledgement

Research reported in this publication was supported in part by the Minnesota State Office for Higher Education Spinal Cord Injury and Traumatic Brain Injury Research Grant Program (FP00098975 and FP00093993), by the subsidy allocated to Kazan Federal University for the state assignment in the sphere of scientific activities (No 17.9783.2017/8.9), and the National Cancer Institute (NCI) of the National Institutes of Health (NIH) under Award Number K99CA214523. The content is solely the responsibility of the authors and does not necessarily represent the official views of the National Institutes of Health.

## Author contributions

S.T., C.C., P.S., S.C., R.I., and I.L. designed the experiment. S.T., C.C., P.S., R.I., P.G., U.L., and I.L. drafted the manuscript. S.T., C.C., P.S., R.I., C.H. collected experiment data. S.T. and P.S. wrote the algorithms for data processing. C.C., R.I., H.W., B.E.K. conducted the animal surgeries. All authors reviewed and participated in editing the manuscript.

## SI Appendix – Video captions

**Supplemental Video 1.** Hemodynamic response with sub-threshold intensity

**Supplemental Video 2.** Hemodynamic response - below fUS cutoff

**Supplemental Video 3.** Hemodynamic response - above fUS cutof

## References

1. Capogrosso, M., et al. (2013) A computational model for epidural electrical stimulation of spinal sensorimotor circuits. J Neurosci 33(49): p. 19326–40.

2. Kapural, L. (2014) Spinal Cord Stimulation for Intractable Chronic Pain. Current Pain and Headache Reports 18(4): p. 406.

3. Young, R.F. and S.J. Stanley (1979) Dorsal Spinal Cord Stimulation in the Treatment of Multiple Sclerosis. Neurosurgery 5(2): p. 225–230.

4. Agari, T. and I. Date (2012) Spinal Cord Stimulation for the Treatment of Abnormal Posture and Gait Disorder in Patients With Parkinson’s Disease. Neurologia medico-chirurgica 52(7): p. 470–474.

5. Fénelon, G., et al. (2012) Spinal cord stimulation for chronic pain improved motor function in a patient with Parkinson’s disease. Parkinsonism & Related Disorders 18(2): p. 213–214.

6. Harkema, S., et al. (2011) Effect of epidural stimulation of the lumbosacral spinal cord on voluntary movement, standing, and assisted stepping after motor complete paraplegia: a case study. The Lancet 377(9781): p. 1938–1947.

7. Angeli, C.A., et al. (2014) Altering spinal cord excitability enables voluntary movements after chronic complete paralysis in humans. Brain 137(5): p. 1394–1409.

8. Harkema, S.J., et al. (2018) Normalization of Blood Pressure With Spinal Cord Epidural Stimulation After Severe Spinal Cord Injury. Frontiers in Human Neuroscience 12(83).

9. Grahn, P.J. et al. (2017) Enabling Task-Specific Volitional Motor Functions via Spinal Cord Neuromodulation in a Human With Paraplegia. Mayo Clinic Proceedings 92: 544–554.

10. Gill M.L., et al. (2018) Neuromodulation of lumbosacral spinal networks enables independent stepping after complete paraplegia. Nat Med. 24(11): 1677–1682

11. Ladenbauer, J., et al. (2010) Stimulation of the human lumbar spinal cord with implanted and surface electrodes: a computer simulation study. IEEE Trans Neural Syst Rehabil Eng 18(6): p. 637–45.

12. Rattay, F., K. Minassian, and M.R. Dimitrijevic (2000) Epidural electrical stimulation of posterior structures of the human lumbosacral cord: 2. quantitative analysis by computer modeling. Spinal Cord 38: p. 473.

13. Lavrov, I., et al. (2006) Plasticity of spinal cord reflexes after a complete transection in adult rats: relationship to stepping ability. J Neurophysiol 96(4): p. 1699–710.

14. Gerasimenko, Y.P., et al. (2006) Spinal cord reflexes induced by epidural spinal cord stimulation in normal awake rats. J Neurosci Methods 157(2): p. 253–63.

15. Lavrov, I. and J. Cheng (2008) Methodological optimization of applying neuroactive agents for the study of locomotor-like activity in the mudpuppies (Necturus maculatus). Journal of Neuroscience Methods 174(1): p. 97–102.

16. Lavrov, I. and J. Cheng (2004) Activation of NMDA receptors is required for the initiation and maintenance of walking-like activity in the mudpuppy (Necturus Maculatus). Canadian Journal of Physiology and Pharmacology 82(8-9): p. 637–644.

17. Cuellar, C.A., et al. (2017) The Role of Functional Neuroanatomy of the Lumbar Spinal Cord in Effect of Epidural Stimulation. Front Neuroanat. 11: 82.

18. Electromyography, in Wiley Encyclopedia of Electrical and Electronics Engineering.

19. Gad P., et al. (2012) Forelimb EMG-based trigger to control an electronic spinal bridge to enable hindlimb stepping after a complete spinal cord lesion in rats. Journal of neuroengineering and rehabilitation 9 (1): 38.

20. Logothetis, N.K., et al. (2001) Neurophysiological investigation of the basis of the fMRI signal. Nature 412: p. 150.

21. Hämäläinen, M., et al. (1993) Magnetoencephalography---theory, instrumentation, and applications to noninvasive studies of the working human brain. Reviews of Modern Physics, 65(2): p. 413–497.

22. Burns, H.D., et al. (2007) MK-9470, a positron emission tomography (PET) tracer for in vivo human PET brain imaging of the cannabinoid-1 receptor. Proceedings of the National Academy of Sciences 104(23): p. 9800–9805.

23. Mace, E., et al., (2011) Functional ultrasound imaging of the brain. Nat Methods 8(8): p. 662–4.

24. Mace, E., et al. (2013) Functional ultrasound imaging of the brain: theory and basic principles. IEEE Trans Ultrason Ferroelectr Freq Control 60(3): p. 492–506.

25. Song P., et al. Functional Ultrasound Imaging of Spinal Cord Hemodynamic Responses to Epidural Electrical Stimulation: A Feasibility Study, Frontiers in Neurology 10: p1–13.

26. Deffieux, T., et al. (2018) Functional ultrasound neuroimaging: a review of the preclinical and clinical state of the art. Current Opinion in Neurobiology 50: p. 128–135.

27. Urban, A., et al. (2015) Real-time imaging of brain activity in freely moving rats using functional ultrasound. Nature Methods 12: p. 873.

28. Lavrov, I., et al. (2008) Facilitation of Stepping with Epidural Stimulation in Spinal Rats: Role of Sensory Input. The Journal of Neuroscience 28(31): p. 7774–7780.

29. Lavrov, I., et al. (2008) Epidural Stimulation Induced Modulation of Spinal Locomotor Networks in Adult Spinal Rats. The Journal of Neuroscience 28(23): p. 6022–6029.

30. Lavrov, I., et al. (2015) Integrating multiple sensory systems to modulate neural networks controlling posture. Journal of Neurophysiology 114(6): p. 3306–3314.

31. Lavrov, I., et al. (2015) Activation of spinal locomotor circuits in the decerebrated cat by spinal epidural and/or intraspinal electrical stimulation. Brain Research 1600: p. 84–92.

32. Gad, P., et al. (2013) Neuromodulation of motor-evoked potentials during stepping in spinal rats. Journal of Neurophysiology 110(6): p. 1311–1322.

33. Shah, P.K. and I. Lavrov (2017) Spinal Epidural Stimulation Strategies: Clinical Implications of Locomotor Studies in Spinal Rats. The Neuroscientist 23(6): p. 664–680.

34. Foroosh, H., J.B. Zerubia, and M. Berthod (2002) Extension of phase correlation to subpixel registration. IEEE Transactions on Image Processing 11(3): p. 188–200.

35. Yu, A.C.H. and L. Lovstakken (2010) Eigen-based clutter filter design for ultrasound color flow imaging: a review. IEEE Transactions on Ultrasonics, Ferroelectrics, and Frequency Control 57(5): p. 1096–1111.

36. Urban, A., et al. (2014) Chronic assessment of cerebral hemodynamics during rat forepaw electrical stimulation using functional ultrasound imaging. NeuroImage, 2014. 101: p. 138–149.

37. Siclari, F., et al. (2007) Developmental anatomy of the distal vertebral artery in relationship to variants of the posterior and lateral spinal arterial systems. AJNR Am J Neuroradiol 28(6): p. 1185–90.

38. Romanes, G.J. (1965) The arterial blood supply of the human spinal cord. Paraplegia 2: p. 199.

39. Herren, R. and L. Alexander (1939) Sulcal and intrinsic blood vessels of human spinal cord. Archives of Neurology & Psychiatry 41(4): p. 678–687.

40. Anderson, N.E. and E.W. Willoughby (1987) Infarction of the conus medullaris. Annals of Neurology 21(5): p. 470–474.

41. Bowen, B.C. (1999) MR Angiography of Spinal Vascular Disease: What about Normal Vessels? American Journal of Neuroradiology 20(10): p. 1773–1774.

42. Huber L, Ivanov D, Handwerker DA, Marrett S, Guidi M, Uludag K, et al. (2018) Techniques for blood volume fMRI with VASO: from lowresolution mapping towards sub-millimeter layer-dependent applications. NeuroImage. 164:131–43.

43. Kemper VG, De Martino F, Emmerling TC, Yacoub E, Goebel R. (2018) High resolution data analysis strategies for mesoscale human functional MRI at 7 and 9.4T. NeuroImage. 164:48–58.

44. Nielsen, A.N. and M. Lauritzen (2001) Coupling and uncoupling of activity-dependent increases of neuronal activity and blood flow in rat somatosensory cortex. The Journal of Physiology 533(3): p. 773–785.

45. Gerasimenko, I., et al. (2001) Initiation of locomotor activity in spinalized cats by epidural stimulation of the spinal cord. Rossiiskii fiziologicheskii zhurnal imeni I.M. Sechenova 87(9): p. 1161–1170.

46. Dickenson, A.H. and D’Mello R. (2008) Spinal cord mechanisms of pain. BJA: British Journal of Anaesthesia, 101(1): p. 8–16.

47. Malisza, K.L. and Stroman P.W. (2002) Functional imaging of the rat cervical spinal cord. Journal of Magnetic Resonance Imaging, 16(5): p. 553–558.

48. Jeon, Y.H. (2012) Spinal cord stimulation in pain management: a review. The Korean journal of pain 25(3): p. 143–150.

49. Errico, C., et al. (2016) Transcranial functional ultrasound imaging of the brain using microbubble-enhanced ultrasensitive Doppler. NeuroImage 124(Pt A): p. 752–761.

50. Seidel, G. and Meairs S. (2009) Ultrasound Contrast Agents in Ischemic Stroke. Cerebrovascular Diseases, 27(suppl 2)(Suppl. 2): p. 25–39.

51. Deng, L., et al., (2016) A multi-frequency sparse hemispherical ultrasound phased array for microbubble-mediated transcranial therapy and simultaneous cavitation mapping. Physics in medicine and biology, 61(24): p. 8476–8501.

