## Supplement material for "Changes in spinal cord hemodynamic reflect modulation of spinal network with different parameters of epidural stimulation"

### Supplemental Materials

##### *Cutoffs for fUS and EMG*

A cutoff was set to distinguish a case group with silent EMG by subjective determination based on visual inspection to EMG signals. As shown in Fig. S1(a), the cutoff for ΔEMG is determined to be 8%, above which the EMG signal can be distinguished from the background activity. Similarly, a cutoff of ΔSCBV was also determined. In Fig. S1(b), where a fUS response with a peak ΔSCBV equal to 2.2% is showed, the response curve fluctuates after the EES ends and does not go back to the baseline. The fluctuation amplitude is larger than half of the peak response. This fluctuation can also be observed from the corresponding hemodynamic response video (Supplemental video 2). In contrast, fUS response with a peak ΔSCBV equal to 3.2%, as shown in Fig. S1(c) (and Supplemental video 3), is a reasonable response curve for the parameter measurement. Therefore, in this study, the cutoff for fUS response was set to be 3%.

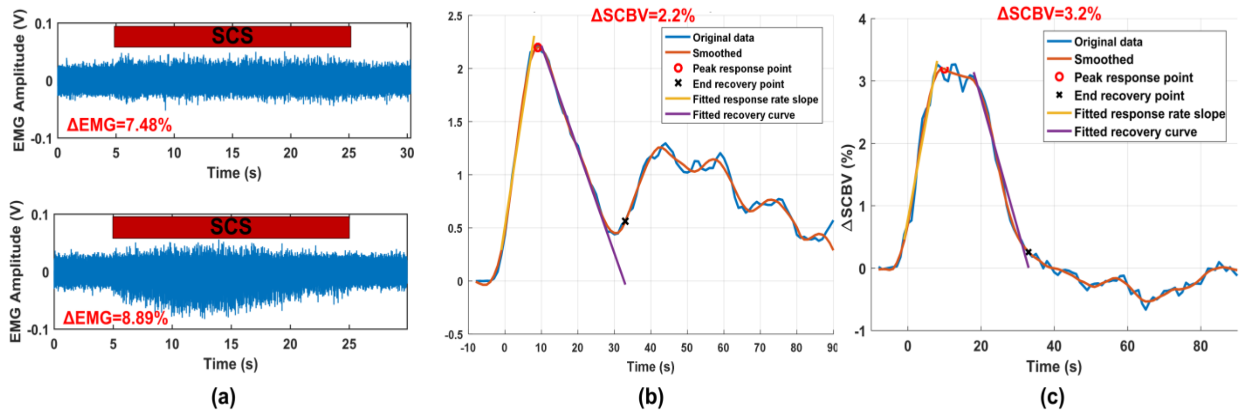


Fig. S1. Cutoff determination for ΔEMG and ΔSCBV. (a) ΔEMG cutoff was set to be 8%, above which the EMG signal can be distinguished from the background activity. (b) ΔSCBV response lower than the cutoff (3%) fluctuates after EES ends with amplitude larger than half of the peak response. (c) ΔSCBV response higher than the cutoff shows a reasonable response curve for the parameter measurement.
